# Conserved gene expression plasticity in development is more pervasive than expression divergence between species of *Caenorhabditis* nematodes

**DOI:** 10.1101/2025.08.04.668491

**Authors:** Athmaja Viswanath, Daniel D. Fusca, John A. Calarco, Asher D. Cutter

## Abstract

Diverse regulatory mechanisms enable precise spatio-temporal control of gene expression across developmental stages, tissues, and sexes, contributing to the proper development of the organism. Evolutionary divergence leads to species-specific gene expression patterns, even in preserved developmental structures, due to regulatory changes that can disproportionately influence subsets of developmental genetic networks. Here we quantify the evolution of sex-biased and tissue-biased transcriptomes from two tissue types (gonad and soma) for each of two sexes (male and female) from two of the closest known sister species of *Caenorhabditis* nematodes (*C. remanei* and *C. latens*). Differential gene expression and co-expression network analyses identify gene sets with distinct transcriptomic profiles, revealing widespread divergence between these morphologically and developmentally cryptic sister species. The transcriptomic divergence occurs despite most genes showing conserved expression across tissues and sexes. These observations implicate shared selection pressures related to tissue and sex differences as outweighing species-specific selection and developmental system drift in shaping overall transcriptome profiles. Although developmentally-plastic tissue-biased expression profiles are mostly conserved between species, we find that sex-biased genes, particularly male-biased genes, contribute disproportionately to species-differences in gene expression, consistent with a disproportionate role for male-biased selection driving gene expression divergence.

## Introduction

Dynamic gene regulation is essential for organismal development and homeostasis, and its evolution is also crucial for adaptation (Carroll, 2000; Prud’homme et al., 2007; Stern, 2000; Wray, 2007; Wray et al., 2003). Gene regulatory elements—such as *cis*-regulatory sequences located near focal genes and *trans*-acting factors encoded elsewhere in the genome—interact with one another and with other gene products to shape the structure and function of gene regulatory networks (GRNs) (Halfon, 2017; Mack & Nachman, 2017). GRNs often are composed of modular sub-networks that work in a synchronized manner to perform distinct functions (Halfon, 2017), allowing for adaptively plastic control over gene expression to mediate conflicting selection pressures across different cell types, developmental stages, and sexes (Breschi et al., 2016; Cutter, 2023a; Cutter & Bundus, 2020; Mendelson & Safran, 2021; Romero et al., 2012). Gene regulatory networks are also able to withstand some disruptions to the network through functional redundancy, where redundant transcription factors and enhancers work together to preserve gene expression and prevent dysfunction of developmental programs in organisms (Arcuschin et al., 2023; MacNeil & Walhout, 2011). Nevertheless, independently evolving populations accumulate distinct mutations that can cause divergence in gene regulatory networks. Species-specific selection provides an obvious potential driver of species differences in gene expression profiles (Halfon, 2017; Mack & Nachman, 2017; Romero et al., 2012). In addition, developmental system drift also can allow regulatory divergence to accrue and gene expression profiles to diverge, even when developmental phenotypes remain conserved (Cutter & Bundus, 2020; True & Haag, 2001). Notable examples of regulatory divergence impacting key developmental processes include the evolution of the Hox transcription factors playing an important role in establishing body plans of vertebrates (Belting et al., 1998; Carroll, 2000) and differential expression of BMP4 and Calmodulin influencing the beak morphology of Darwin’s finches (Abzhanov et al. 2004, 2006; Wu et al. 2004). Consequently, it remains important to assess how divergence in gene regulatory control over sex and tissue development may intersect with evolutionary forces to generate conserved versus species-specific patterns throughout the transcriptome.

The modular and partially-independent nature of GRNs enables the possibility of differential evolution of transcriptome profiles among developmentally-distinct cell types (i.e. tissue-biased expression due to differential regulation) (Breschi et al., 2016; Cutter & Bundus, 2020; Mendelson & Safran, 2021; Romero et al., 2012). Among animal tissues, brain tissues have shown the highest conservation (Brawand et al., 2011; Khaitovich et al., 2005), while genes expressed in the testis exhibit substantial divergence in expression between species (Brawand et al., 2011). Moreover, genes with narrow, tissue-specific expression patterns tend to evolve more rapidly than do genes with broad expression and more pleiotropic roles (Khaitovich et al., 2005). Studies of the evolution of tissue-biased expression between species of mammals, however, provide evidence for greater variation between distinct tissues of the same species than between homologous tissues across species (Blake et al., 2020; Brawand et al., 2011; Khaitovich et al., 2005; Romero et al., 2012), albeit with some conflicting evidence (Blekhman et al., 2008; Brawand et al., 2011; Lin et al., 2014; Yanai et al., 2004) and some dependence on the particular tissues under consideration (B.-Y. Liao & Zhang, 2006; Romero et al., 2012). While gene expression and gene coding sequence mostly remain conserved between species, different tissues exhibit substantial variation in the rate of gene expression evolution (Brawand et al., 2011; Khaitovich et al., 2005). These observations raise the question of how universal across animals are predispositions of genes expressed in certain tissue types to rapid evolution, and how often are gene expression differences greater between tissues due to developmental plasticity than between species due to evolution.

Much like how gene regulatory mechanisms can cause tissue-biased expression, they can also yield developmentally plastic gene expression between individuals as sex-biased differences in transcriptomes (Tosto et al., 2023). Sex-biased gene expression may arise from quantitative differences or sex-limited expression, leading to distinct genetic network architectures between sexes (Tosto et al., 2023; Wright et al., 2018). Consequently, sex-biased gene expression is likely shaped by both sexual selection and sex-biased natural selection (Tosto et al., 2023), as well as conflicting selective pressures between sexes known as sexual antagonism, such that sex-biased gene expression can help resolve fitness conflicts over sex differences in optimal expression (Ellegren & Parsch 2007; Cutter 2023a). Moreover, male-biased genes tend to possess a greater tissue specificity than sex-neutral genes, implying that mutations will cause fewer pleiotropic effects and so can respond more efficiently to a given selective pressure (Tosto et al., 2023). Sex-biased genes, specifically male-biased genes, are often observed to evolve more rapidly than female-biased and sex-neutral genes both in coding sequence and expression level (Assis et al., 2012; Cutter & Ward, 2005; Ellegren & Parsch, 2007; Khaitovich et al., 2005; Meiklejohn et al., 2003; Reinke et al., 2004). Consequently, genes showing sex-biased expression may contribute disproportionately to transcriptome differences between species (Ellegren & Parsch, 2007; Khaitovich et al., 2005; Meiklejohn et al., 2003; Ranz et al., 2003).

Here, we aimed to test how sex-biased and tissue-biased gene expression contribute to species-specific differences using the *Caenorhabditis* nematode sister species *C. remanei* and *C. latens* (Dey et al., 2012; Félix et al., 2014). These morphologically and developmentally cryptic species form one of the closest known relatives of *Caenorhabditis* nematodes, having diverged <5 MYA with strong but incomplete post-zygotic reproductive isolation (Bundus et al., 2018; Dall’Acqua et al., 2025; Dey et al., 2012; Félix et al., 2014; Fusca et al., 2025; Memar et al., 2019). Studies of other *Caenorhabditis* species have shown an important role for sex-specific gene expression contributing to differences within and between species (Albritton et al., 2014; Kramer et al., 2016; Sánchez-Ramírez et al., 2021; Thomas et al., 2012), in addition to conservation in transcriptome profiles during embryogenesis (Large et al., 2025; Levin et al., 2012). To gain further evolutionary genetic insight into regulatory divergence of plastic gene expression across developmental contexts, we quantified and analyzed transcriptomes for gonad and somatic tissues for adults of each sex of each species to determine profiles of differential expression both within and between species, for both orthologous and species-specific genes.

## Methods

### Nematode dissection and RNA isolation

We cultured populations of young adult *C. remanei* (VX0003) and *C. latens* (VX0088) on NGM-agar plates with *Escherichia coli* OP50 at 25°C. To collect gonad and somatic tissues, ∼400 young adult worms were dissected per sample using a 0.3mm x 30mm needle (BD # 305109). To isolate the tissues, the worms were initially cut behind the pharynx while in sterile water. This process led to the release of the gonads outside the body of the worm. Subsequently, the gonadal arms were separated from the carcass by making a second cut. We pooled together ∼400 separated gonads to form one “gonad” tissue sample and the rest of the ∼400 carcasses to form one “somatic” tissue sample. Gonad and somatic tissue samples were collected in triplicate separately for males and females of each species. All the samples were stored in Tri-reagent at - 80°C for ∼48-72 hours after which RNA was isolated using Zymo RNA Isolation kit (#R2051). The isolated RNA was then treated with Turbo DNase (#AM2238) and RiboLock RNase Inhibitor (#EO0381) to degrade any DNA in the samples. To get high quality and high yield of RNA, the samples were further purified using Zymo RNA Clean and Concentrator kit (#R1015) according to manufacturer protocols.

### Sequencing and transcript analysis

We generated high-quality mRNA sequence (RNA-seq) data for 24 samples in total, corresponding to 3 replicates for each of 8 treatments (2 Species × 2 Sexes × 2 Tissues). Each sample yielded an average >75 million paired-end reads (read length 150bp) generated using NovaSeq S4 flowcell. The cDNA library preparation and next generation sequencing was performed by The Centre for Applied Genomics, The Hospital for Sick Children, Toronto, Canada. We used updated genome annotations for *C. remanei* PX506 (Teterina et al., 2020) and *C. latens* PX534 v2 (Adams et al., 2023) reference genomes by incorporating existing annotations along with the RNAseq data generated in this study. These updated annotations were generated by the Centre for the Analysis of Genome Evolution and Function, University of Toronto, Toronto, Canada, with the repeat sequences in the genomes were analyzed with RepeatModeler2 using the default parameters (Flynn et al., 2020) (gff annotation files available as Supplementary Files S1 & S2). Sequence reads were quality trimmed with Trimmomatic (Bolger et al., 2014) (parameters: LEADING:3 TRAILING:3 SLIDINGWINDOW:4:15 MINLEN:36), and duplicate reads were removed with Nubeam-dedup using the default parameters (Dai & Guan, 2020). Transcript assembly and annotation were conducted as described in Rifkin et al. (2022). We mapped the quality-verified mRNA reads against the reference genomes and updated annotations of the respective species for each sample using STAR with default parameters (Dobin et al., 2013). We quantified gene expression of all the genes using FeatureCounts (Y. Liao et al., 2014) and restricted subsequent analysis of the read count data to the set of 13,785 1:1 orthologous genes or to species-specific singleton genes identified using Orthofinder (Emms & Kelly, 2015).

### Estimation of Coding Sequence Divergence and Codon Usage Bias

DNA coding sequences for all transcripts from each genome assembly were retrieved using GffRead v0.12.3 (Pertea & Pertea, 2020). For every pair of 1:1 orthologs, the *C. remanei* and *C. latens* coding sequences were aligned to each other using PRANK v.170427 (Löytynoja, 2014) in codon-alignment mode. dN and dS values were estimated for each alignment with the FitMG94 script from HyPhy v2.5.52 (Kosakovsky Pond et al., 2020) after replacing all stop codons with gaps. For each 1:1 ortholog alignment, we also calculated the Effective Number of Codons (ENC) with coRdon v1.14.0 (Elek et al., 2025), using the *C. remanei* sequence from each alignment. We then calculated a corrected measure of dS (dS′) for each 1:1 ortholog alignment, to correct for the effect of codon usage bias on estimates of dS. We used the linear correction of dS′ = dS + 0.006926114 * (61 – ENC), where 0.006926114 is the slope of the linear regression between dS and ENC calculated for these alignments (excluding alignments with dS > 1 from the regression, as these likely represent alignment errors or incorrect orthology inferences), and 61 is the value of ENC for a gene with no codon usage bias.

### Differential gene expression analysis

Differential gene expression analysis was performed for orthologous genes using DESeq2 (Love et al., 2014) in R Studio (RStudio version 4.3.2). In DESeq2 we used the design formula: ∼batch + species + sex + tissue + species: sex + species: tissue + sex: tissue to identify the effects of different factors and their interactions on gene expression. Benjamini-Hochberg false discovery rate-adjusted P_adj_ < 0.05 was used as a criterion for identifying differentially expressed between treatments from DESeq2 contrasts to infer genes with species-biased expression, sex-biased expression, and tissue-biased expression.

### Weighted gene co-expression network analysis

The R package WGCNA was used to perform a Weighted Gene Co-Expression Network Analysis to identify co-expressed modules of genes that were different across species, sexes, and tissues (Langfelder & Horvath, 2008). We performed WGCNA using the combined dataset comprising all 24 samples, resulting in 11 distinct modules representing gene set clusters with highly correlated expression profiles (Langfelder & Horvath, 2008). Differences in filtering criteria led WGCNA to use a slightly smaller gene set than DESeq2 (n_WGCNA_ = 11,804 orthologous genes, n_DESeq2_ = 13,428).

We represented the stereotypical expression profile for each module by a module eigengene, which corresponds to its first principal component in expression space. We also associated module expression profiles with species, sexes and tissues using Pearson’s Correlation Coefficient in WGCNA. Intramodular “hub” genes were inferred using two metrics: gene significance and module membership (Langfelder & Horvath, 2008). Gene significance (GS) correlates the expression of a gene to external trait or condition information which in this case was sex, tissue and species. The higher the absolute value of GS, the more significantly associated that gene is with the trait or condition. GS of zero indicates that the gene is not associated with the trait or condition. Module membership (MM) is the correlation between a given gene’s expression and the module eigengene (first principal component of a module). Highly connected intramodular hub genes show high MM magnitudes (close to +1 or -1). Using these two measures, we classified genes in M10 and M4 with GS >0.9 and MM >0.9 as “hub” genes (Langfelder & Horvath, 2008) that considered to be highly connected with other genes in their module and show significant difference in expression between species.

### Data Availability Statement

The mRNA sequencing reads are available under the accession number GSE304374. All code related to the analyses presented in this study can be found in the following GitHub repository: https://github.com/Athmaja-Viswanath/Evolution-of-expression-plasticity-Caenorhabditis

## Results

### Tissue- and sex-dependent gene expression differences are more prevalent than species divergence in gene expression

To characterize the combined effects of species, sex, and tissue leading to differential gene expression, we performed differential gene expression analysis using DESeq2 (Love et al., 2014). Our analysis design formula included all main effects — species (Sp), sex (S), and tissue (T) — as well as their pairwise interactions (∼Sp + S + T + Sp×T + Sp×S + T×S) for 13,428 orthologous genes with detectable non-zero expression, successfully classifying them into genes with significant differential expression (DEG) and non-DEG. We visualized shared and unique expression patterns for DEGs (Figure 1), excluding the 717 genes (5.3%) with indistinguishable expression across all treatments (non-DEG). Among the 12,711 genes that showed some form of significant differential expression, approximately 70% (n = 9156 genes) showed a simple dependence on tissue, sex, species, or their additive effects (Figure 1). Overall, tissue type and sex exerted a stronger influence on gene expression than species differences (Figure 1, Supplementary Figure S1), with roughly 50% more genes being differentially expressed between tissues or sexes relative to between species. The preponderance of tissue and sex differences over species differences points to within-species regulatory differences — across tissues and sexes — being more prominent than divergence between species. Consequently, expression dissimilarity is generally greater between different tissues or sexes than between species for the same tissue or sex.

**Figure 1:**
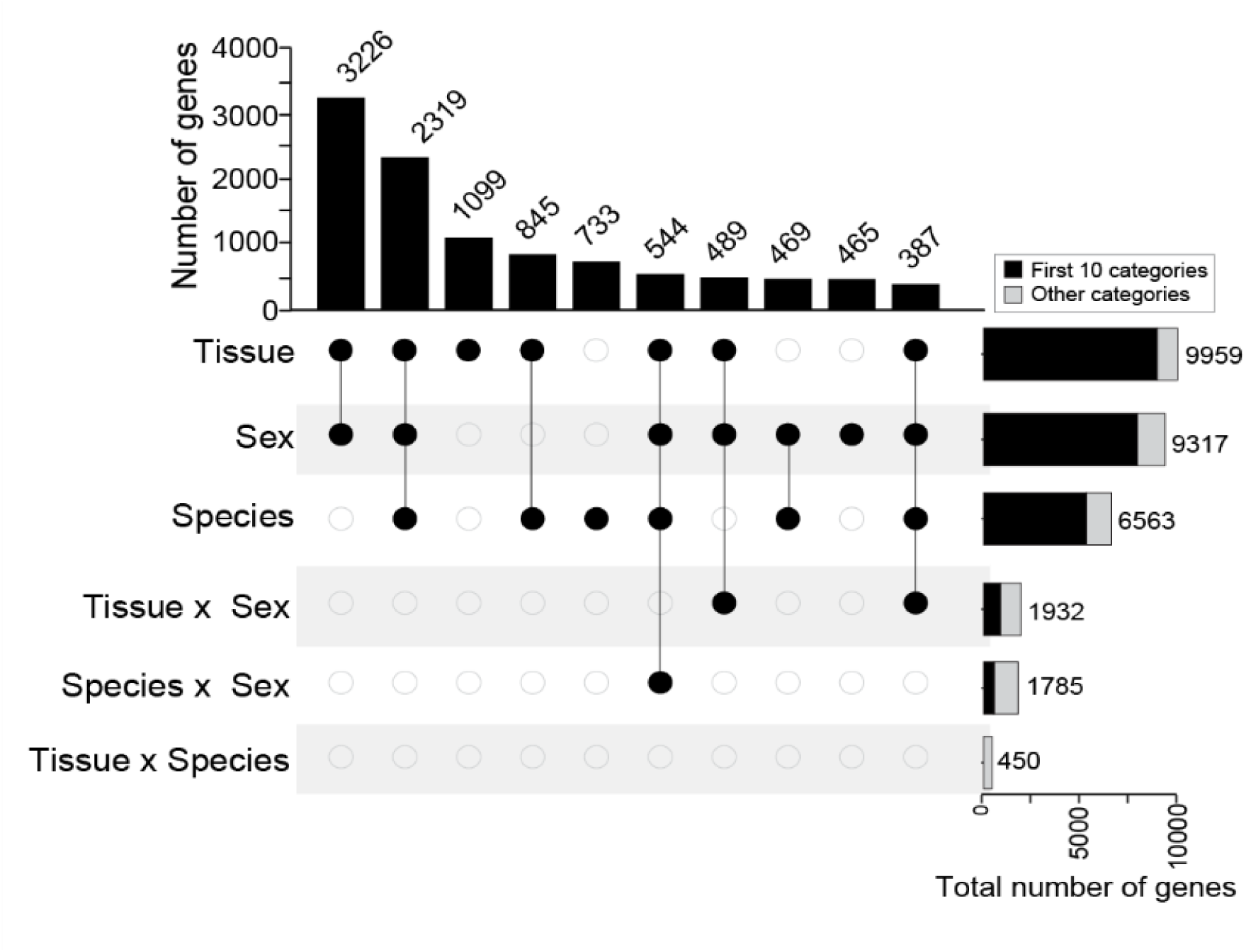
Tissue and sex-differences play a larger role in generating gene expression differences than species-differences. Upset plot depicting the number of genes in the 10 largest gene categories derived from differential gene expression analysis using DESeq2. These 10 categories contribute to ∼83% of total differential gene expression (n= 10,576 genes out of total 12,711 genes across 59 “significance categories”, Supplementary Figure S1); 2135 DEGs corresponding to rarer combinations of significance categories not shown for simplicity. The horizontal bars represent the total number of genes in each category. The 717 non-DEGs are not included.

### Extensive gene expression divergence between *C. remanei* and *C. latens*

Despite the pronounced plasticity of gene expression between tissues and sexes within species, ∼49% of 1:1 gene orthologs between *C. remanei* and *C. latens* nonetheless show differential expression between the species (n_DEG_ = 6563 of 13,428 gene orthologs that were successfully classified by DESeq2), with the remainder showing conserved expression between species (n_Conserved_ = 6865) (Figure 2A, Supplementary Table S1). Nearly equal numbers of genes showed higher expression in each species (n*_Crem_* = 3224, n*_Clat_* = 3339). However, species-biased genes showed the lowest density on the X-chromosome (44% of assayed X-linked genes) and highest density on chromosome II (∼52%) (Figure 2B, Figure 5C). Functional enrichment analysis using gProfiler (Kolberg et al., 2023) identified species-biased genes related to pathways involving reproduction, negative regulation of cellular processes, response to stimulus, catalytic activity, among others (Figure 2C). Genes showing species-biased expression also exhibited significantly higher rates of coding sequence divergence compared to genes with conserved expression, as measured by dN/dS ratios (dN/dS for *Crem*-biased vs. conserved GLM, t = -11.493, P < 0.0001; *Clat*-biased vs. conserved GLM, t = 8.306, P < 0.0001) (Figure 5B, Supplementary Table S1). Moreover, we also observed a positive correlation between the magnitude of expression divergence and coding sequence divergence (dN/dS) across species-biased genes (Supplementary Figure S3). The abundance of differentially expressed genes (DEG) between species demonstrates that substantial regulatory divergence has accumulated between the genomes of *C. remanei* and *C. latens* over the several million years since they shared a common ancestor, despite being morphologically and developmentally cryptic species (Dey et al., 2014).

**Figure 2:**
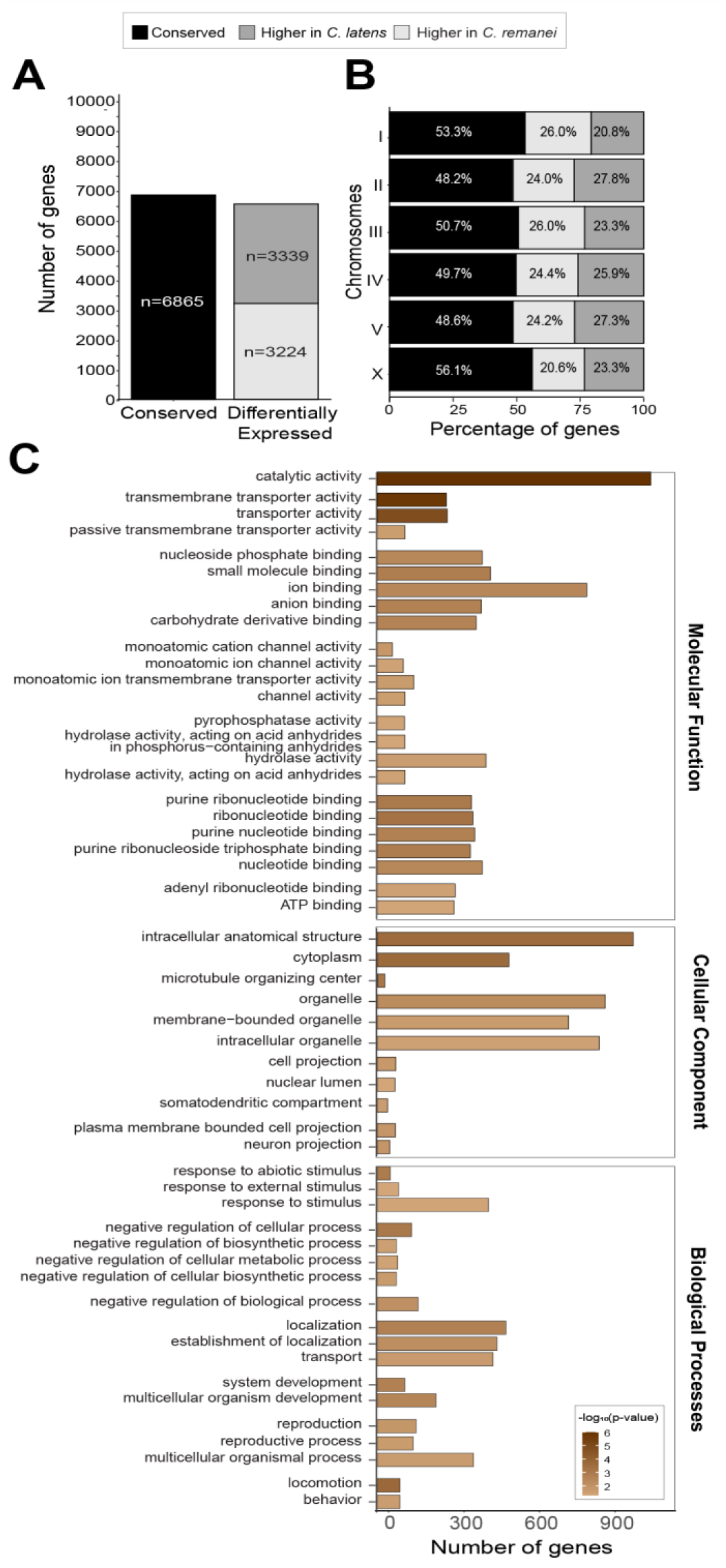
Differential gene expression analysis reveals substantial gene expression divergence between *C. remanei* and *C. latens*. (A) Bar plot depicting the number of conserved genes (black) and *C. remanei*-biased genes (light grey) and *C. latens*-biased genes (dark grey). (**B**) Chromosomal distribution of orthologous genes across *C. remanei* chromosomes. (**C**) Functional enrichment analysis of *C. remanei* biased genes using gProfiler. Only significant GO terms are shown. *C. latens*-biased genes did not show enrichment of any term.

### Regulatory divergence may predominate in species differences, especially among X-linked genes

We next sought to understand gene co-expression patterns jointly between tissues and sexes in the context of divergent expression between species with weighted gene co-expression analysis (WGCNA) (Langfelder & Horvath, 2008). We identified 11 distinct gene co-expression modules (M1-M11) that detailed sets of genes with similar expression profiles among orthologs (Figure 3A), corroborating disproportionate influence of within-species plasticity of expression by tissue and sex relative to between-species differences. Six modules were significantly associated with only one factor (species, tissue, or sex), two correlated with both tissue and sex, and the remaining three showed expression differences between species and either tissue or sex (Figure 3A, B). The four largest modules accounted for nearly 75% of all genes (27% n_M5_ = 3156, 23% n_M9_ = 2761, 15% n_M2_ = 1772, 8% n_M10_ = 979 out of 11804 genes).

**Figure 3:**
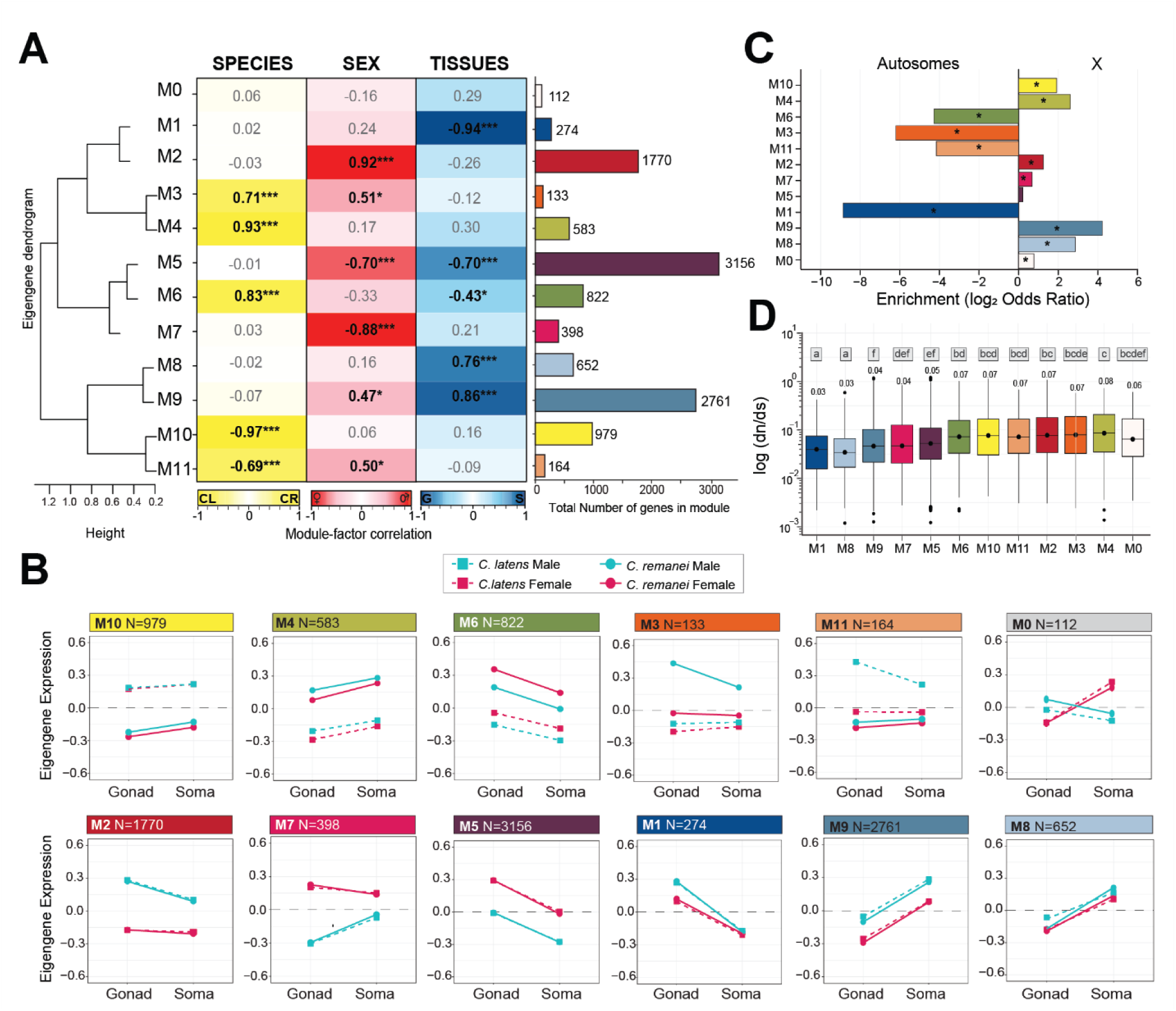
WGCNA identified 11 co-expression modules correlated with stronger expression by species, sex, and tissue. (**A**) Module-factor correlation plot indicates Pearson’s *r* correlation coefficient as calculated between module eigengene (first principal component of a module) and each factor (species, sex, tissue) to identify modules correlated with species, sex, and tissue. Module M10, M2 and M1 are most significantly correlated with species, sex, and tissue respectively and are depicted by primary colors (M10 *r* = -0.97 (yellow), M2 *r* = 0.92 (red), M1 *r* = -0.94 (dark blue)). All other modules are represented by combinations of these three primary colours to depict the relative effect of different conditions on module eigengene expression. Genes that could not be clustered in any of the 11 modules were assigned to Module 0 (M0). The eigengene dendrogram (left) represents the similarity between different modules. The barplot (right) indicates the total number of genes in each module, with M5 and M9 being the largest modules (n_M5_ = 3156, n_M9_ = 2761). (**B**) Module eigengene expression patterns across species, sexes, and tissues. The blue lines correspond to males and red lines indicate females (solid for *C. remanei*, dashed for *C. latens*). (**C**) Enrichment of genes on autosomes versus X-chromosomes for different modules as log_2_(odds ratio). A positive value indicates enrichment on X-chromosome and a negative value indicates enrichment on autosomes (Fisher’s exact test; * indicates significant enrichment P < 0.05). (**D**) Median dN/dS values for genes across different modules (interquartile range and 95^th^ percentiles indicated by box and whiskers); numerical median dN/dS value indicated above each module. Letters above each boxplot indicate statistical difference categories according to pairwise Tukey’s post-hoc tests; modules sharing the same letter are not significantly different (P > 0.05), while those with different letters show significant differences (P < 0.05).

Two modules (M4 and M10) were significantly associated with species identity alone and represent sets of genes with global differences between species and which, interestingly, display a significant >2-fold enrichment on the X-chromosome to implicate a disproportionate role for X-linked gene regulation in overall species-biased gene expression divergence (Figure 3C). We identified 271 (M10) and 134 (M4) highly connected “hub genes” in these two modules (Figure 4A, C). Hub genes—those showing high connectivity within a module—are expected to play key roles in gene regulation and essential biological processes due to their extensive interactions within co-expression networks (Langfelder & Horvath, 2008; F.-L. Zhang et al., 2023). However, rates of coding sequence divergence were similar for both hub and non-hub genes (M10 GLM, t = 1.614, P = 0.1069; M4 GLM, t = -1.518, P = 0.1295) (Figure 4B, D; Supplementary Table S8), despite hub genes being enriched in M10 for roles in the formation of protein complexes, including ribosome and ribonucleoprotein complexes. More generally, genes in those expression modules that correlated significantly with species identity generally did not exhibit significantly elevated rates of coding sequence evolution compared to modules associated with sex biased expression, though dN/dS ratios were roughly two-fold higher relative to modules associated most strongly with tissue biased expression (Figure 3D; Supplementary Table S9). Together, these findings further reinforce the idea that species differences may largely arise from regulatory divergence.

**Figure 4:**
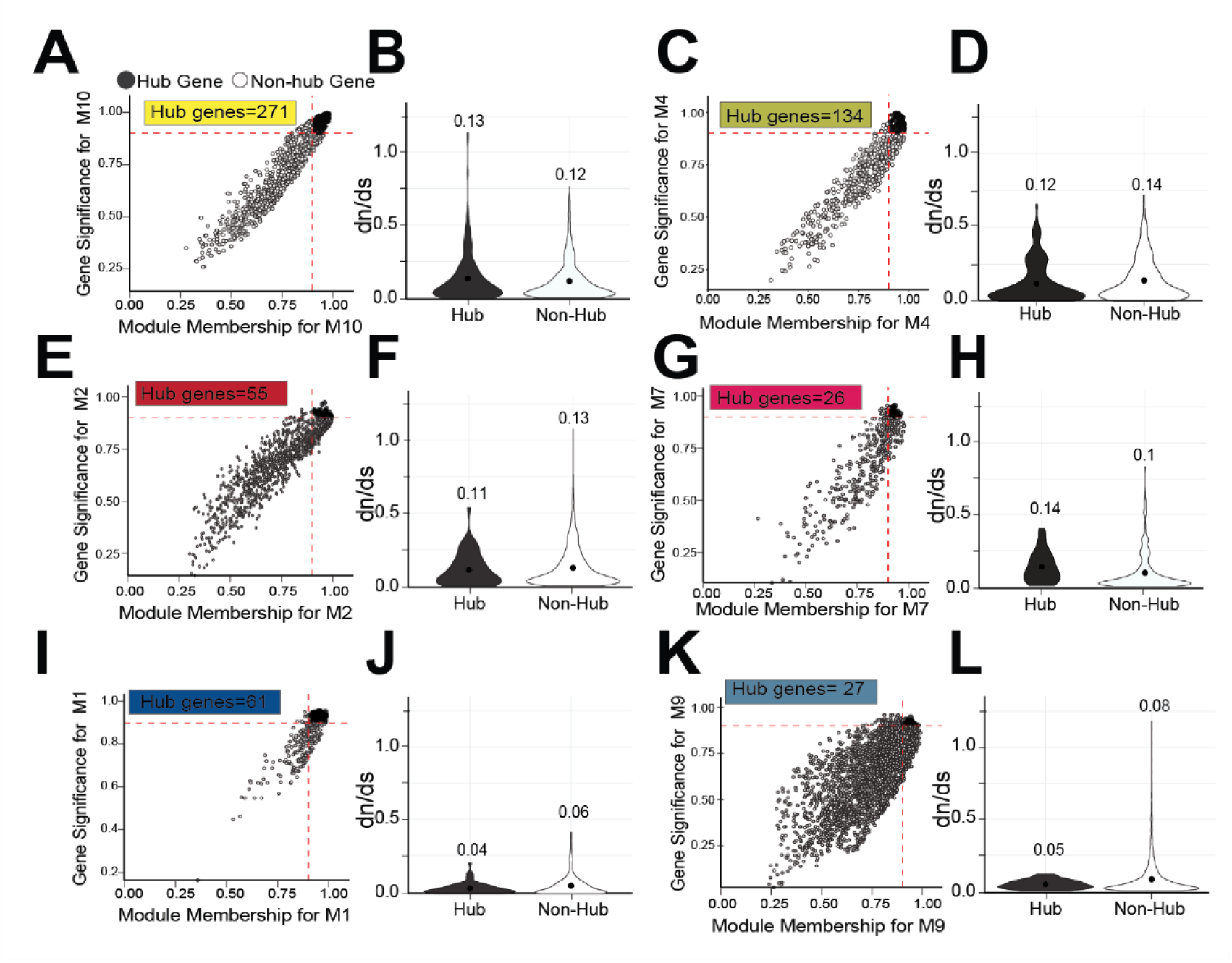
Tissue-biased modules display significantly different rates of molecular evolution between hub and non-hub genes. Scatter plots (**A, C, E, G, I, K**) illustrating “hub” genes (black points) in species-biased modules M10 (**A**) and M4 (**C**), sex-biased modules M2 (**E**) and M7 (**G**), and tissue-biased modules M1 (**I**) and M9 (**K**). Corresponding violin plots (**B, D, F, H, J, L**) comapre dN/dS distribution between hub genes (black) and non-hub genes (white). No significant differences were observed for species-biased modules M10 (**B**; GLM, t = 1.614, P = 0.1069) and M4 (**D**; t = -1.518, P = 0.1295), or sex-biased modules M2 (**F**; GLM, t = -0.772, P = 0.4403) and M7 (**H**; GLM, t = 1.283, P = 0.2004). In contrast tissue-biased modules M1 (**J**; GLM, t = **-**2.191, P = 0.0293) and M9 (**L**; GLM, t = -2.159, P = 0.0309) show slower dN/dS in hub genes suggesting stronger evolutionary constraint.

### Disproportionate male-biased gene expression in sex-bias and species differences

Male-biased expression defined four of six co-expression modules inferred by WGCNA to significantly correlate with sex, comprising ∼35% more genes (M2, M3, M9, M11; n=4828) than the two female-biased co-expression modules (M5, M7; n=3554) (Figure 3A). Notably, male-biased modules M3 and M11, enriched >4-fold on autosomes, also correlated with species, suggesting divergent sex-biased regulation between *C. remanei* and *C. latens* (Figure 3C) and consistent with a role for male-biased selection in driving divergence between species in gene expression. In contrast, two other modules showed sex differences with no significant correlation with either species identity or tissue (male-biased M2, female-biased M7), consistent with these sets of genes having largely conserved sex-biased expression across tissues and species. Genes in M2 and M7 were also enriched on the X-chromosome, implicating conserved X-linkage of some sources of sex-biased gene expression for both sexes (Figure 3C, Supplementary Table S8). The rates of coding sequence divergence were similar for hub and non-hub genes in sex-biased modules M2 (Figure 3E, F; GLM, t = -0.772, P = 0.4403) and M7 (Figure 4G, H; GLM, t = 1.283, P = 0.2004).

Consistent with the findings with WGCNA co-expression modules, differential gene expression analysis revealed that genes with male-biased expression were more prevalent than genes with either female-biased or sex-neutral expression (38.6%, n_male-biased_ = 5184; 31%, n_female-biased_ = 4133; 30.4%, n_sex-neutral_ = 4111 out of 13,428 genes). Male-biased genes are disproportionately abundant on autosomes in contrast to sex-neutral genes which are enriched on the X-chromosome (Figure 5C, Supplementary Figure S2), consistent with prior findings (Albritton et al., 2014; Reinke et al., 2004). Male-biased genes displayed higher average expression levels than female-biased and sex-neutral genes (male-biased vs. female-biased GLM, t =-26.830, P < 0.0001; |log2-fold-difference|, male-biased vs. sex-neutral GLM, t = -70.275, P <0.0001, female-biased vs. sex-neutral GLM, t = 41.352, P <0.0001) (Figure 5A, Supplementary Table S2). Coding sequence evolution is faster for male-biased genes only when compared to female-biased genes (dN/dS GLM, t =-7.184, P < 0.0001), though genes with sex-neutral expression showed slightly elevated coding sequence evolution compared to genes with male-biased expression (dN/dS GLM, t = 2.760, P = 0.0160) (Figure 5B, Supplementary Table S2). The faster molecular evolution of male-biased genes compared to female-biased genes is consistent with some predictions of differential sexual selection pressures potentially contributing to species-differences (Dapper & Wade, 2020; Ellegren & Parsch, 2007; Wade, 2002).

**Figure 5:**
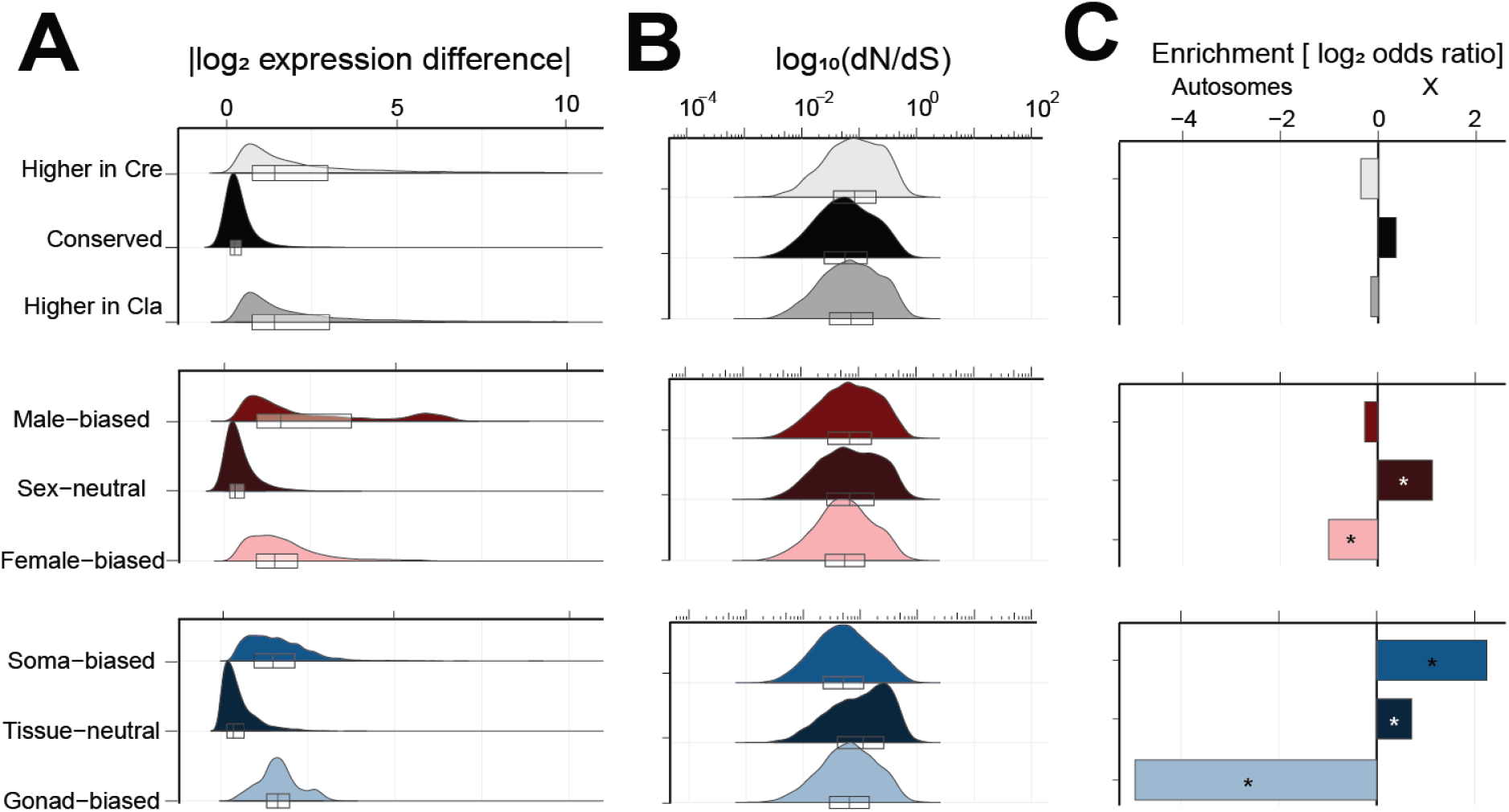
Male-biased genes display higher expression difference and sequence divergence. (**A**) Absolute expression difference |log2-fold-difference| and (**B**) coding sequence divergence (dN/dS) are compared across species-, sex- and tissue-biased genes. Male-biased genes display higher expression difference than female-biased (GLM, *t* =-26.830, P < 0.0001) and sex-neutral genes (GLM, *t* = -70.275, P <0.0001); female-biased genes also differ from sex-neutral genes (GLM, t = 41.352, P <0.0001). Coding sequence evolution is faster for male-biased genes only when compared to female-biased genes (dN/dS GLM, t =-7.184, P < 0.0001), though genes with sex-neutral expression showed slightly elevated coding sequence evolution compared to genes with male-biased expression (dN/dS GLM, t = 2.760, P = 0.0160). Tissue-biased genes showed slower coding sequence evolution compared to tissue-neutral genes (tissue-neutral vs. gonad-biased, t = - 17.716, P < 0.0001; tissue-neutral vs. soma-biased, t =22.403, P < 0.0001; gonad-biased vs. soma-biased, t =6.069, P < 0.0001) but displayed higher expression difference (gonad-biased vs. tissue-neutral, t = 77.135, P <0.0001; soma-biased vs. tissue-neutral, t = - 72.308, P < 0.0001). (**C**) Chromosomal enrichment analysis. Species-biased genes are not significantly enriched on autosomes or X-chromosome (n_soma-biased_ = 4472, 33.6%; n_gonad-biased_ = 5437, 40.9%; n_tissue-neutral_ = 3383, 25.5% of 13,292 genes); Fisher’s exact test, P < 0.05 and |log2 odds ratio < 0.5|. In contrast, sex-neutral genes are enriched on the X-chromosome while female-biased genes were autosome-enriched (Fisher’s exact test; * indicates significant enrichment P < 0.05). Gonad-biased genes show strong autosomal enrichment (>4- fold) and are more abundant than soma-biased or tissue-neutral genes (n_soma-biased_ = 4494, 33.5%, n_gonad-biased_ = 5465, 40.7%; n_tissue-neutral_ = 3469, 25.8% of 13,428 genes), whereas tissue-neutral and soma-biased genes showed X-chromosome enrichment (Fisher’s exact test; * indicates significant enrichment P < 0.05).

To further explore sex-biased gene expression in species differences, we examined species-specific genes of *C. remanei* and *C. latens* (single-copy non-orthologous genes; n*_Crem_* = 982 genes, n*_Clat_* = 2279 genes). We found sex-biased species-specific genes of both species to be much rarer than sex-neutral genes when compared to their relative incidences among 1:1 orthologs (n*_Crem_*_, male-biased_ = 152, 25.8%; n *_Crem_*_, female-biased_ = 136, 23.1%; n *_Crem_*_, sex-neutral_ = 301, 51.1% of 589 total genes that were categorized by DESeq2) (n*_Clat_*_, male-biased_ = 327, 27.3%; n *_Clat_*_, female-biased_ = 185, 15.4%; n *_Clat_*_, sex-neutral_ = 685, 57.4% of 1197 genes categorized by DESeq2) (Figure 6B, Supplementary Table S3 & S4). Similar to orthologs, however, species-specific male-biased genes displayed higher average expression levels compared to sex-neutral genes and female-biased genes (*C. remanei* |log2-fold-difference|, male-biased vs. sex-neutral GLM, t = - 31.492, P <0.0001; male-biased vs. female-biased GLM, t =-8.691, P < 0.0001, Supplementary Table S3; *C. latens* |log2-fold-difference|, male-biased vs. sex-neutral GLM, t = -31.492, P <0.0001; male-biased vs. female-biased GLM, t =-8.691, P < 0.0001, Supplementary Table S4) (Figure 6A, B).

**Figure 6:**
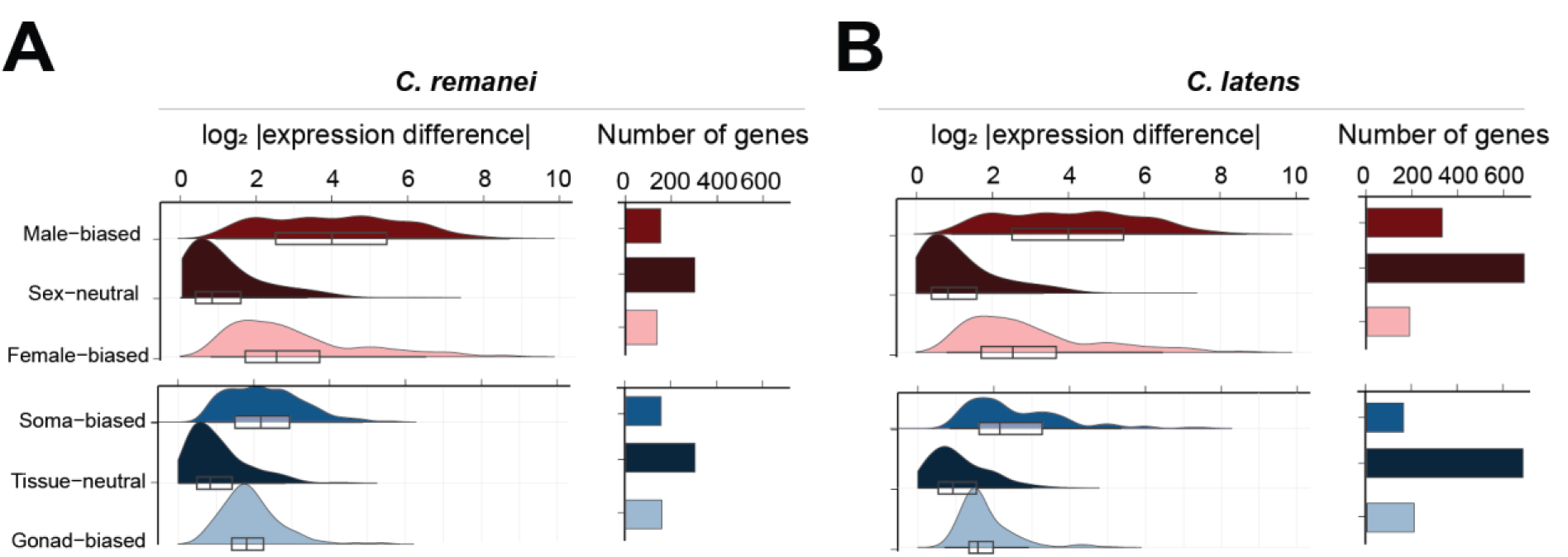
Male-biased and soma-biased genes display greater differential expression among species-specific genes. Distributions of the magnitude of differential expression among treatment categories for non-orthologous genes between sexes and tissues in (**A**) *C. remanei* (n_Crem, male-biased_ = 152, 25.8%; n _Crem, female-biased_ = 136, 23.1%; n _Crem, sex-neutral_ = 301, 51.1% of 589 total genes; |log2-fold-difference|, male-biased vs. sex-neutral GLM, t = -31.492, P <0.0001; male-biased vs. female-biased GLM, t =-8.691, P < 0.0001; gonad-biased vs. tissue-neutral GLM, t = 10.526, P <0.0001; soma-biased vs. tissue-neutral GLM, t =- 15.187, P < 0.0001; soma-biased vs. gonad-biased GLM t= -4.130, P=0.0001) and (**B**) *C. latens* (n_Clat, male-biased_ = 327, 27.3%; n _Clat, female-biased_ = 185, 15.4%; n _Clat, sex-neutral_ = 685, 57.4% of 1197 genes; |log2-fold-difference|, male-biased vs. sex-neutral GLM, t = -31.492, P <0.0001; male-biased vs. female-biased GLM, t =-8.691, P < 0.0001; gonad-biased vs. tissue-neutral GLM, t = 10.067, P <0.0001; soma-biased vs. tissue-neutral GLM, t = -19.356, P <0.0001; soma-biased vs. gonad-biased GLM, t =-8.545, P < 0.0001). Barplots indicate the number of genes in each gene category.

### High conservation of expression and sequence among tissue-biased genes

We next considered how tissue-biased (gonad and soma) expression can intersect with species differences. For orthologous genes, five co-expression modules showed tissue-biased profiles of expression, with 24.6% more genes represented among the three modules indicating gonad-biased expression (M1, M5, M6, n= 4252) than for somatic-biased expression (M8, M9, n=3413 genes) (Figure 3A). Only one module also associated significantly with species differences (M6: higher expression in *C. remanei* and in gonad tissue), consistent with strong conservation of expression for most genes with tissue-biased expression. Interestingly, 2 of the 3 gonad-biased modules show gene enrichment on autosomes whereas the soma-biased modules are enriched on the X-chromosome (Figure 3C), consistent with sensitivity to sex chromosome linkage in tissue-biased gene expression (Reinke et al., 2000, 2004). Average rates of coding sequence divergence are consistently lower for genes in tissue-biased modules than for genes in sex-biased and species-biased modules (Figure 3D, Supplementary Table S5). Hub genes of tissue-biased modules show more conserved coding sequence evolution than non-hub genes (dN/dS, M1 GLM, t = -2.191, P = 0.0293; M9 GLM, t = -2.159, P = 0.0309) (Figure 4I, J, K, L), in contrast to our observations for hub gene status in modules exhibiting sex-bias and species-biased expression.

Differential expression analysis with DESeq2 corroborated these WGCNA observations about chromosomal enrichments and sequence conservation (Figure 5). Genes with gonad-biased expression were >4-fold enriched on autosomes (Figure 5C, Supplementary Figure S2) and more abundant than either soma-biased or tissue-neutral genes (n_soma-biased_ = 4494, 33.5%, n_gonad-biased_ = 5465, 40.7%; n_tissue-neutral_ = 3469, 25.8% of 13428 genes). Tissue-biased genes as a whole displayed higher average expression levels compared to tissue-neutral genes (|log2-fold-difference|, gonad-biased vs. tissue-neutral GLM, t = 77.135, P <0.0001; soma-biased vs. tissue-neutral GLM, t = - 72.308, P < 0.0001) (Figure 5A). We also observed that genes with tissue-biased expression showed slower coding sequence evolution compared to tissue-neutral genes (dN/dS, tissue-neutral vs. gonad-biased, GLM, t = - 17.716, P < 0.0001; tissue-neutral vs. soma-biased GLM, t =22.403, P < 0.0001) (Figure 5B, Supplementary Table S5). Together, these patterns implicate strong purifying selection for tissue-biased genes in both expression and coding sequences to yield few changes to the gonad versus somatic expression status of genes between sexes and species. Nonetheless, gonad-biased genes displayed faster coding sequence evolution than soma-biased genes (gonad-biased vs. soma-biased, GLM, t = 6.069, P < 0.0001) (Figure 5B, Supplementary Table S5). Consequently, tissue-dependent expression imposes distinctive evolutionary constraints, with gonad-biased genes evolving more rapidly due to either elevated positive selection or less potent negative purifying selection.

When we separately performed differential expression analysis for genes unique to the genomes of either *C. remanei* or *C. latens* (Figure 6), we found tissue-biased genes to be unusually rare compared to what we observed for 1:1 orthologs (*C. remanei* n_gonad-biased_ = 158, 25.6%; n_soma-biased_ = 155, 25.2%; n_tissue-neutral_ = 303, 49.2% of 616 genes) (*C. latens* n_gonad-biased_ = 205, 19.6%; n_soma-biased_ = 159, 15.2%; n_tissue-neutral_ = 680, 65.1% of 1044 genes) (Figure 6B). Among the species-specific tissue-biased genes that we could analyze, however, they displayed higher average expression level compared to tissue-neutral genes, a pattern similar to what we found for 1:1 orthologs (*C. remanei* |log2-fold-difference|, gonad-biased vs. tissue-neutral GLM, t = 10.526, P <0.0001; soma-biased vs. tissue-neutral GLM, t =-15.187, P < 0.0001; soma-biased vs. gonad-biased GLM t= -4.130, P=0.0001, Supplementary Table S6; *C. latens* |log2-fold-difference|, gonad-biased vs. tissue-neutral GLM, t = 10.067, P <0.0001; soma-biased vs. tissue-neutral GLM, t = -19.356, P <0.0001; soma-biased vs. gonad-biased GLM, t =-8.545, P < 0.0001, Supplementary Table S7) (Figure 6A, B). These observations reinforce the notion of weaker conservation for tissue-unbiased gene expression that may also manifest in terms of more pervasive gene turnover as represented by non-ortholog species-specific genes.

## Discussion

Biologically distinct species of *Caenorhabditis* nematodes show exceptional conservation in outward morphology and ontogenetic cell lineage development (Memar et al., 2019), raising the question of how conserved their transcriptomes are in the face of extensive genome sequence change (Large et al., 2025). We demonstrated that sister species *C. remanei* and *C. latens* have diverged substantially in transcriptome profiles in constitutive as well as tissue-specific and sex-specific ways. Despite being one of the closest known pairs of *Caenorhabditis* species and capable of hybridization, their distinct gene expression profiles involve ∼49% of orthologous genes in species-biased expression. Nonetheless, their transcriptome divergence is less extreme than seen for another close pair of *Caenorhabditis* species (54%-60% expression divergence between *C. briggsae* and *C. nigoni*) (Sánchez-Ramírez et al., 2021), which also have diverged starkly in reproductive mode, genome size, and gene composition (Ren et al., 2018; Yin et al., 2018). These observations reinforce the conclusion that divergence between even closely-related species in transcriptome profiles can be substantial, as documented in flies (Ranz et al., 2003; S. A. Rifkin et al., 2003), mice (Kautt et al. 2024), and plants (Davidson et al., 2012). Unlike some systems, however, genes showing differential expression between *C. remanei* and *C. latens* did not cluster along chromosomes in a manner consistent with “islands of speciation” in the genome (Mendelson & Safran, 2021; Ravinet et al., 2017). Moreover, despite the widespread transcriptome divergence between *C. remanei* and *C. latens*, we found that plastic differential expression due to tissue identity and sex was even more pronounced. These findings reinforce the conclusions from single-cell transcriptomics across embryogenesis for distantly-related *C. elegans* and *C. briggsae* that nonetheless document extensive conservation in gene expression, despite divergence (Large et al., 2025). That is, different tissues or sexes of the same species generally show greater expression dissimilarity than do distinct species for the same tissue or sex.

### Gene expression conservation and divergence in the evolution of developmental genetic networks

Differences in gene expression between species can be driven by the divergence of gene regulatory pathways that govern development (Kalinka et al. 2010; Levin et al. 2012; Hashimshony et al. 2015, 2016; Memar et al. 2019; Cutter et al. 2019; Cutter & Bundus 2020). An important question to answer, then, is how gene regulation evolves under tissue-specific selective pressures to cause divergence in developmental gene regulatory networks. In line with earlier findings (Blake et al., 2020), we rarely observe tissue-biased modules exhibiting species-specific differences in orthologous genes, indicative of conserved tissue-biased gene expression between species. Furthermore, we found that genes in tissue-biased modules consistently exhibit lower average rates of coding sequence divergence compared to those in sex-biased and species-biased modules. Additionally, within tissue-biased modules, hub genes show more conserved coding sequence evolution than non-hub genes—contrasting with the pattern observed in sex-biased and species-biased modules.

Taken together, these observations support the notion that stabilizing expression is especially strong on this class of genes, with gene expression more likely to be similar between homologous tissues of different species than between different tissues within the same species (B.-Y. Liao & Zhang, 2006; Romero et al., 2012). While stabilizing selection to maintain gene expression profiles can act at the sequence level by weeding out mutations, it also manifests as the fixation of compensatory mutational changes in *cis-* and *trans*-acting factors such that they preserve gene expression profiles between species (Mack & Nachman, 2017; Signor & Nuzhdin, 2018). In this way, gene expression as a trait can be conserved across species, despite differences in the genetic encoding of regulatory mechanisms (Gilad et al., 2006; Lemos et al., 2005; Mack & Nachman, 2017). Consequently, our analysis leaves open the question of whether such conservation in expression emerges due to conserved regulation or, instead, due to divergent but compensatory regulatory mechanisms that serve to maintain overall gene expression profiles, as a form of developmental system drift (DSD) (Cutter, 2023b; Cutter & Bundus, 2020). Conservation of expression, as for soma-biased genes, could also imply that the soma-biased gene regulatory networks remain robust despite mutational changes and thus maintain the same phenotype.

We identified only one tissue-biased module that also showed differential expression between species (M6, enriched for gonad-biased genes), which also exhibited faster coding sequence evolution among its genes than for modules containing soma-biased genes. Both of these observations point to an important contribution of reproductive evolution to species differences. Interestingly, such differences between species could manifest as dysfunctional developmental programs in interspecies hybrids (Civetta & Singh, 1995). For instance, *C. remanei* and *C. latens* produce viable hybrids of both sexes, but hybrid males show gonad dysfunction and sterility in one direction of the cross (Dey et al., 2014). We consider this issue further, below, with respect to sex-biased expression divergence.

Apart from transcriptional regulation involving *cis*-regulatory elements, other regulatory mechanisms also can contribute to species-differences in observed gene expression. Gene duplication, gain, or loss events can impact the divergence of gene regulation among different species, affecting transcriptome expression by altering gene dosage or neofunctionalization leading to distinct gene regulatory networks between species (Lynch & Force, 2000; Panchy et al., 2016; Taylor & Raes, 2004). In addition to shared orthologs, we examined the expression of genes that are unique to one species or the other, which also may contribute importantly to gene regulatory networks controlling developmental programs. While we found that genes showing tissue-biased expression among species-specific genes were unusually rare, they might nonetheless lead to novel developmental gene regulatory network architecture in non-germline and germline tissues, contributing to species-specific differences (Lynch & Force, 2000).

Moreover, species-specific differences in gene composition can potentially lead to divergence in developmental networks between species to contribute to reproductive isolation (Halfon, 2017; S. A. Rifkin et al., 2003; Wagner & Misof, 1993). For example, many duplicated genes have male-biased expression in *Drosophila*, including *OdsH*, a duplicate of *unc-4* that evolved a new function contributing to hybrid incompatibility in flies (Ting et al., 2004); gain-of-function activity from F-box gene family expansion in *C. nigoni* also contributes to genetic incompatibility with its sister species *C. briggsae* (Xie et al., 2024). It remains to be determined what the relative importance is for changes to ortholog gene expression versus gene copy number to overall divergence between species in transcriptome profiles and genetic incompatibility between species.

Gene expression evolution also can be affected by divergence in alternative splicing, small RNA regulatory pathways, and mediators of epigenetic modification to contribute to species differences in transcriptome profiles (Blake et al., 2020; Merkin et al., 2012; Miller & Matute, 2016; Patlar et al., 2019; Rogers et al., 2021). The identity of tissue being compared also is crucial to understanding the evolution of tissue-biased regulation, as differences in cell type compositions even within similar tissues can lead to different gene expression patterns (Church et al., 2023; Hunnicutt et al., 2022; Price et al., 2022). Since we considered all non-gonad tissue as somatic tissue, this composite set of ∼1000 somatic cells per animal comprises multiple tissue types (e.g. muscle, intestine, neuron, etc.) and may have limited our ability to detect more subtle patterns of tissue-biased gene expression divergence among distinct cell types between the species. Future exploration of the full spectrum of gene regulatory mechanisms and specific cell types is crucial for gaining a comprehensive understanding of how genetic networks evolve across species.

### The role of sex-biased gene expression in creating species differences

Sex-biased gene expression is a common outcome of gene regulatory evolution across different taxa, though its extent depends on the tissue and developmental stage of the organism (Grath & Parsch, 2016). Sex-biased gene regulation is important as it can help resolve intra-locus sexual conflict where different sexes have different fitness optima for a shared trait, such as the amount or timing of gene expression in development (Cutter, 2023a; Tosto et al., 2023; Wright et al., 2018). Sex-biased regulatory divergence also can drive novel gene activity to influence key biological processes like reproduction (Assis et al., 2012). Our findings confirm that male-biased genes, in particular, are major contributors to species differences in expression. Only those sex-biased co-expression modules that exhibited a male expression bias showed significant associations with species-differences for orthologous genes. Additionally, genes specific to either *C. remanei* or *C. latens* that were male-biased in expression displayed higher average expression levels compared to sex-neutral genes and female-biased genes. This disproportionate contribution of male-biased genes to divergence between species in their transcriptome profiles reinforces findings about sex-biased gene expression evolution in other systems (Assis et al., 2012; Llopart, 2012; Meiklejohn et al., 2003; Patlar & Civetta, 2021; Ranz et al., 2003; Sánchez-Ramírez et al., 2021).

Sex-biased genes, particularly male-biased genes, also often evolve rapidly in protein sequence (Assis et al., 2012; Grath & Parsch, 2016), as seen in mammals, insects, and other nematodes (Cutter & Ward, 2005; Torgerson et al., 2002; Z. Zhang et al., 2004). Differences in protein evolution between male- and female-biased genes can be attributed to male-biased genes experiencing disproportionate positive selection due to sexual selection or sex-biased directional natural selection and/or female-biased genes facing stronger evolutionary constraints due to more pleiotropic roles (Assis et al., 2012; Grath & Parsch, 2016; Mendelson & Safran, 2021; Parisi et al., 2004), especially when associated with gonad expression. For instance, positive selection can drive divergence in seminal fluid protein coding genes (Patlar & Civetta, 2021); spermatogenesis-related genes exhibit higher protein sequence divergence in many taxa, often associated with antagonistic coevolution with egg recognition proteins and inter-male competition (Howard, 1999; Levitan & Stapper, 2010; Palumbi, 2009; Vacquier & Swanson, 2011). Alternatively, weaker purifying selection on sex-biased genes might also contribute to the rapid evolution of reproductive genes (Dapper & Wade, 2020). While our results show statistical support for such patterns in *C. remanei* and *C. latens*, the magnitudes of effect were modest for higher rates of protein evolution for genes with male-versus female-biased expression, and did not differ from genes with sex-neutral expression. These findings are thus consistent with a more pronounced influence of divergence in gene regulation than protein sequence in contributing to sex-biased evolution in these species.

These findings underscore the importance of gene regulatory evolution in species divergence, particularly among male-biased genes. They also align with previous reports of high levels of interspecific expression divergence in male-biased genes being driven by changes in gene regulatory elements such as *cis*-acting elements and *trans*-acting factors (Assis et al., 2012; Khaitovich et al., 2005; Llopart, 2012; Mack & Nachman, 2017; Meiklejohn et al., 2003; Ranz et al., 2003). Although the importance of male-biased gene expression in species-level differences is well-known across different taxa, theoretical models demonstrate that sexual selection alone often is insufficient to lead to the completion of speciation (Mack & Nachman, 2017; Mendelson & Safran, 2021; Servedio & Bürger, 2014). Nonetheless, genetic changes favored by sexual selection might contribute to post-zygotic developmental genetic incompatibilities in interspecies hybrid offspring, beyond simple effects of sexual selection on mating and mate choice. It will thus be valuable to determine the molecular genetic causes, in terms of gene regulatory mechanisms, that are responsible for sex-biased gene expression divergence as may be inferred through allele-specific expression analysis in hybrids (Sánchez-Ramírez et al., 2021; Wittkopp et al., 2004). Further work will help to elucidate the nuanced relationship between sexual selection and speciation (Lindsay et al., 2019; Mendelson & Safran, 2021) and, more broadly, how genomes may reveal predictable rules of gene regulatory evolution that underpin expression conservation and differentiation among tissues, sexes, and species.

## Conclusion

We discovered significant differences in gene expression within and between the transcriptomes of nematode sister species *Caenorhabdits remanei* and *C. latens* in a sex- and tissue-dependent manner. Through analyses of differential expression and gene co-expression networks, we demonstrated that multiple selective pressures influence the transcriptomic profiles of divergence and conservation. Our findings implicate shared selection pressures of tissues and sexes among species to be more pronounced than species-specific selection in shaping the overall transcriptome profiles. Despite pervasive conservation of expression profiles, it remains to be determined whether the genomic encoding of regulatory controls of gene expression similarly reflect pervasive conservation in regulatory sequences or, instead, compensatory coevolutionary change of cis- and trans-acting regulators. Genes with male-biased expression, however, we found to contribute disproportionately to species differences in gene expression, consistent with an outsized role for male-biased selection driving gene expression divergence. These findings highlight the intricate interplay of evolutionary forces shaping gene expression profiles within and among species.

## Acknowledgements

We thank Katja Kasimatis and Rebecca Schalkowski for their valuable input on experimental design and RNA extractions, as well as for their helpful comments on data interpretation.

