## Supplementary Information for "Conserved gene expression plasticity in development is more pervasive than expression divergence between species of *Caenorhabditis* nematodes"

### Supplementary Figures

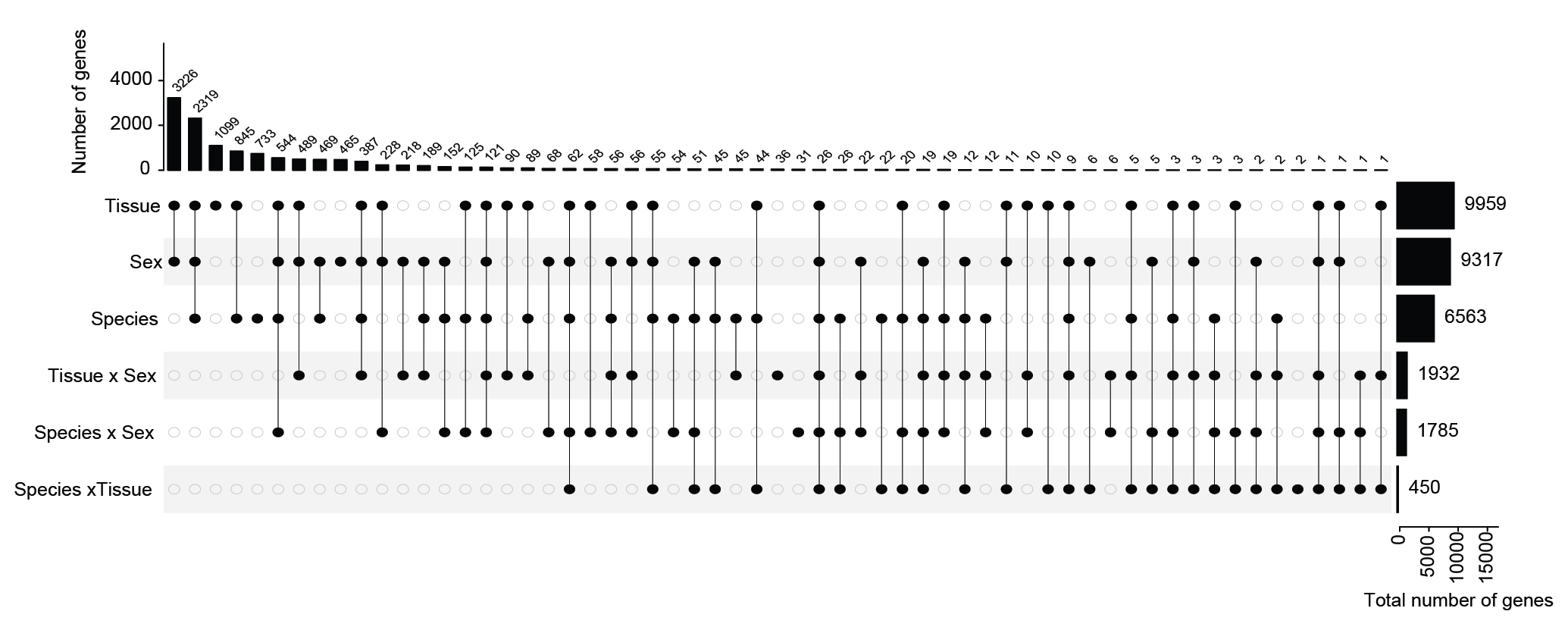

Supplementary Figure S1. Upset plot depicting 12,711 genes across all the 59 “significance categories” from differential gene expression analysis using DESeq2. The 10 largest categories contribute to ~83% of total differential gene expression (n= 10,576 genes out of total 12,711 genes). Horizontal bars represent the total number of genes in each category. The 717 non-DEGs are not included.

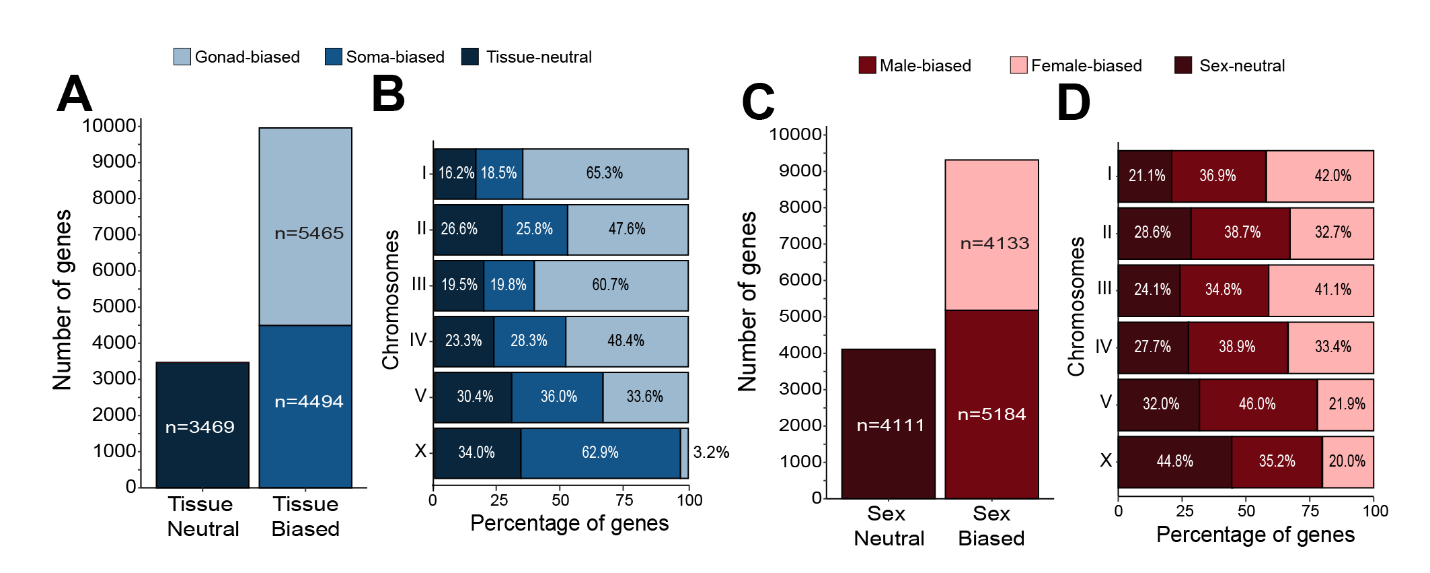

**Supplementary Figure S2:** Distribution of tissue-biased and sex-biased genes across chromosomes. (**A, C**). Overall number of genes displaying tissue-biased (**A**) and sex-biased gene expression (**C**). (**B, C**) Chromosomal distribution of tissue-biased (**B**) and sex-biased genes (**C**).

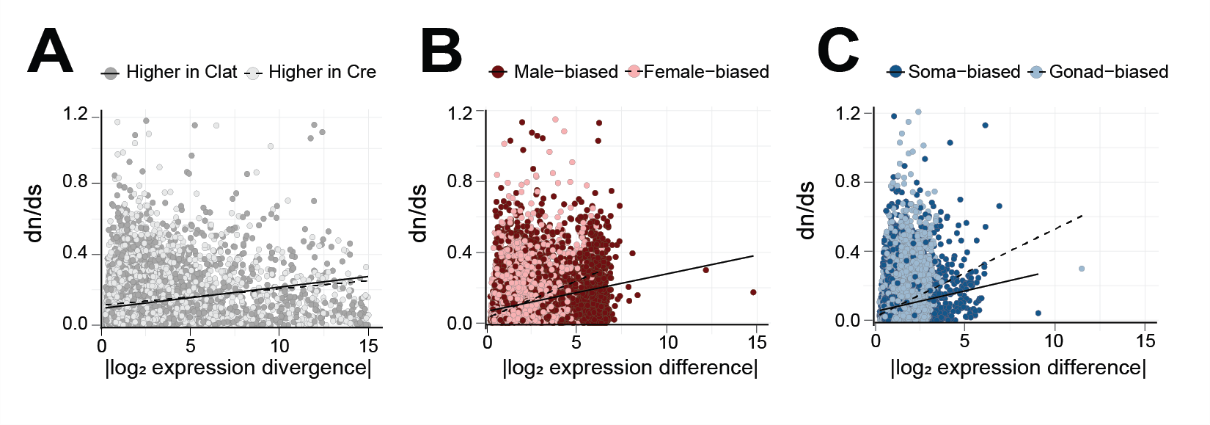

**Supplementary Figure S3**. Biplots depicting the Pearson’s correlation between the magnitude of gene expression (x-axis) and coding sequence divergence dN/dS (y-axis) for: (**A**) species-biased genes (r*_Crem_* = 0.127, *p* <0.001; r*_Clat_*= 0.174, *p* <0.001), (**B**) sex-biased genes (r*_male-biased_* = 0.306, *p* <0.001; r*_female-biased_*= 0.374, *p* <0.001), and (**C**) tissue-biased genes (r*_gonad-biased_* = 0.246, *p* <0.001, r*_soma-biased_* = 0.183, *p* <0.001).

### Supplementary Tables

**Supplementary File S1:** Updated gene annotations for *C. latens*

**Supplementary File S2:** Updated gene annotations for *C. remanei*

**Supplementary Table S1:** Species-biased orthologous genes between *C. remanei* and *C. latens*. The "log2 fold change" column represents the expression difference between *C. remanei* and *C. latens*, with positive values indicating higher expression in *C. remanei*. The "basemean" column shows the average normalized expression across all samples for each gene, adjusted for library size. The “crexclat” column indicates expression bias: 0 = conserved expression, 1 = higher expression in *C. remanei* and -1 = higher expression in *C. latens*. Synonymous substitution rates (dS) are corrected for codon usage bias using effective number of codons metric (ENC) (see dS_corrected).

**Supplementary Table S2:** Sex-biased orthologous genes. "log2 fold change" represents differential expression between males and females, with positive values indicating higher expression in males. The "basemean" indicates average normalized expression. The “mxf” column indicates expression bias: 0 = sex-neutral expression, 1 = male-biased expression and -1 = female-biased expression. dS_corrected values are corrected for codon usage.

**Supplementary Table S3:** Sex-biased *C. remanei*-specific genes. As in Table S2, "log2 fold change" reflects male vs. female expression, and “mxf” indicates expression bias. Data are limited to genes with no identifiable orthologs in *C. latens*.

**Supplementary Table S4:** Sex-biased *C. latens*-specific genes. Same metrics and bias definitions as Table S3, applied to *C. latens*-specific genes.

**Supplementary Table S5:** Tissue-biased orthologous genes. "Log2 fold change" reflects soma vs. gonad expression, with positive values indicating soma-biased expression. "Basemean" is average normalized expression. The gxs column denotes expression bias: 0 = tissue-neutral, 1 = soma-biased, -1 = gonad-biased. Synonymous substitution rates are codon-corrected (dS_corrected).

**Supplementary Table S6:** Tissue-biased *C. remanei*-specific genes. Same metrics and interpretation as Table S5, limited to genes without orthologs in *C. latens*.

**Supplementary Table S7:** Tissue-biased *C. latens*-specific genes. Same format and interpretation as Table S6, focused on *C. latens*-specific genes.

**Supplementary Table S8:** Orthologous genes in WGCNA modules. Includes module assignments and codon-corrected dS values (dS_corrected).

**Supplementary Table S9:** Post-hoc comparison of module eigengenes using Tukey's HSD. Includes adjusted p-values and confidence intervals for post-hoc comparisons between co-expression modules based on eigengene expression.
